# FYCO1 improves postischemic cardiac remodeling via enhanced autophagic flux and attenuation of proinflammatory signaling

**DOI:** 10.64898/2026.03.17.712297

**Authors:** Frauke Senger, Susanne S Hille, Anca Kliesow Remes, Tapan K Baral, Abel Martin-Garrido, Nesrin Schmiedel, Christian Kuhn, Oliver J Müller, Ashraf Y Rangrez, Johannes Backs, Arica Beisaw, Jörg Heineke, Norbert Frey

## Abstract

Acute myocardial infarction (MI) is associated with severe metabolic and oxidative stress that triggers cardiomyocyte death, pro-inflammatory signaling and progressive structural remodeling frequently culminating in heart failure. Although significant advances in reperfusion therapy improved acute survival in patients, therapeutic strategies that directly target intracellular processes in response to injury remain limited. One key response mechanism, autophagy, is rapidly activated to ameliorate ischemic stress. Yet, defective autophagic flux may exacerbate cardiomyocyte injury and maladaptive tissue remodeling. Here we identify FYCO1 as a cardiomyocyte-enriched key regulator of autophagy that enhances autophagic flux and promotes myocardial resilience following ischemic injury. Using cardiomyocyte-specific FYCO1 transgenic mice subjected to permanent coronary ligation, we demonstrate that FYCO1 overexpression limits infarct expansion, reduces cardiomyocyte injury, and preserves cardiac function during remodeling. In vivo RFP-EGFP-LC3 autophagy reporter analyses reveal that FYCO1 promotes a sustained increase of autophagic flux by coordinating autophagosome formation and efficient autolysosomal clearance. Transcriptomic profiling identifies a cardioprotective gene program in FYCO1-Tg animals subjected to MI, with suppression of proinflammatory, proapoptotic and stress-response pathways. Systemic serum cytokine and chemokine profiling as well as transcriptomic analyses of myocardium confirm reduced inflammatory signaling and subsequent reduction in macrophage recruitment into the infarct border zone. Together these findings position FYCO1 as a key regulator of cardiomyocyte autophagy and reveal a previously unrecognized link between autophagy and inflammation in shaping cardiac remodeling following myocardial infarction. FYCO1-mediated autophagy promotes myocardial preservation and functional recovery, highlighting autophagic flux as a promising target for cardioprotective interventions.

## Introduction

Ischemic heart disease remains the leading cause of death worldwide [1–3]. Acute myocardial infarction is defined as acute myocardial injury occurring in the setting of ischemia which initiates a cascade of cellular injury and maladaptive cardiac remodeling [4,5]. Despite significant therapeutic advances, AMI remains a leading cause of global morbidity and mortality, accounting for more than 7 million events annually worldwide; with mortality rates being especially high in low-and middle-income regions [1,6,7]. The growing prevalence of diabetes, obesity, and population aging is expected to further expand the global disease burden [2,8], underscoring the need for improved prevention and therapy [1]. Following coronary artery occlusion, cardiomyocytes under ischemic stress experience severe metabolic and oxidative stress that triggers cell death, pro-inflammatory signaling and progressive structural remodeling advancing into adverse cardiac remodeling [4,5,9,10]. Although significant advances in reperfusion therapy improved acute survival in patients [5], therapeutic strategies that directly target intracellular processes in response to injury remain limited. Understanding the molecular mechanisms as well as their crosstalk that defines the cardiac microenvironment after injury represents a critical step toward developing new cardioprotective interventions [4,11].

Autophagy has emerged as a pivotal cellular quality control mechanism that plays a central role in cardiomyocyte stress adaptation [12–14]. Autophagy is a key adaptive response to myocardial ischemia, acting as an intracellular quality control system that removes damaged proteins and organelles when oxygen and nutrient supply are compromised [12,15–17]. In patients with coronary artery disease and acute myocardial infarction, dynamic changes in autophagy markers, consistent with an ischemia-induced increase in autophagosome formation and context-dependent alterations in autophagic flux were observed [13,15,18]. Several human studies report altered expression of LC3, Beclin-1, ATG5 and p62 in ischemic and failing hearts compared with non-ischemic controls [12,13,18–20], indicating that basal autophagy becomes insufficient or dysregulated as ischemia progresses to chronic adverse remodeling and subsequent heart failure [12,15,19,21].

However, increasing evidence implies that dysregulation of autophagic flux—rather than mere induction of autophagosome formation—contributes to myocardial injury and progression towards heart failure [12,14,15,22]. Inefficient autophagic processing and impaired autolysosomal clearance can lead to accumulation of dysfunctional proteins and organelles, resulting in cardiomyocyte death and proinflammatory signaling during post-infarction remodeling [15]. Enhancing autophagic flux—rather than only increasing autophagosome formation—appears to be a decisive protective mechanism in ischemic human myocardium [12,20,22–24]. Cardiac biopsies from ischemic hearts show upregulation of LC3-II and Beclin-1 and, in many cases, accumulation of p62, which is compatible with enhanced autophagosome formation but impaired lysosomal clearance—a pattern frequently reported in AMI [13,15,18,19]. Preservation of lysosomal function and coordinated autolysosomal clearance have been shown to be essential for cardioprotection [12,22,23].

Several cardioprotective interventions tested in clinical or translational settings— including ischemic conditioning strategies [12], GLP-1 analogues [25], metformin [20,23], and AMPK activators [24]—converge on enhancing autophagic flux rather than simply stimulating autophagy initiation [12,24]. This flux-centric protection likely explains why therapies that inhibit lysosomal degradation, such as chloroquine derivatives [12,26], may worsen cardiomyopathy in exposed patients: blocking autophagolysosomal clearance traps cardiomyocytes in a state of energetic insufficiency and proteotoxic stress [12,26]. Collectively, the existing human and translational evidence indicates that boosting autophagic flux may be a therapeutic opportunity to limit infarct size, preserve viable myocardium, and improve post-ischemic recovery [12,20,23,24,27].

Beyond its canonical degradative role to maintain intracellular homeostasis, autophagy has recently also been recognized as an important regulator of cardiac inflammatory and apoptotic signaling following injury [12,28]. Following MI, necrotic cardiomyocytes release danger-associated molecular patterns [29–32] that trigger cytokine and chemokine secretion [33–36] and initiate rapid on-site recruitment of immune cells into the infarct border zone [31,33,37,38,39]. This early phase inflammatory response is essential for removal of necrotic debris thereby initiating tissue repair [31,40]. Along this line, excessive or prolonged immune activation promotes cardiomyocyte cell death, infarct expansion, and adverse remodeling [11,12,31,32,34,40–43]. However, the molecular mechanisms that coordinate autophagic activity with inflammatory remodeling remain incompletely understood. [12,28,44].

FYCO1 is a cardiac enriched regulator of the autophagic machinery that localizes to late autophagosomal compartments and facilitate their trafficking, thereby enhancing autophagic flux. Previous studies from our group demonstrated that cardiomyocyte-specific overexpression of FYCO1 enhances autophagic flux and preserves cardiac function in heart failure due to pressure overload [45]. Nevertheless, the underlying mechanisms by which FYCO1-driven autophagy may modulate myocardial injury, inflammatory and apoptotic signaling and post-infarction remodeling upon ischemia remain unknown.

Our present study thus investigates the impact of FYCO1-mediated autophagy enhancement in myocardial remodeling following ischemic injury. Using a permanent coronary ligation model combined with in vivo a RFP-EGFP-LC3 in vivo autophagy reporter, transcriptomic profiling and immune phenotyping, we demonstrate that FYCO1 promotes sustained and balanced autophagic flux in cardiomyocytes, limits inflammatory signaling and consequently attenuates immune cell recruitment into the infarct border zone. This coordinated orchestration reshapes the post-infarction microenvironment, resulting in attenuated apoptotic cell death signaling, reduced infarct expansion and preservation of cardiac function. Together, our findings position FYCO1 as a key regulator of cardiomyocyte autophagy and reveal a previously unrecognized link between autophagy and inflammation in shaping remodeling following myocardial infarction.

## Methods

### Animals and Ethical Approval

All animal procedures complied with the institutional and national Guidelines for the Care and Use of Laboratory Animals and were approved by the Ministry of Agriculture, Rural Areas, European Affairs and Consumer Protection of Schleswig-Holstein (Germany) ([MELLUR ID- V 242/ - 13649/2018 (32-4/18)]) and were performed according to EU Directive 2010/63/EU and ARRIVE-guidelines. Double-transgenic mice cardiomyocyte-specific FYCO1-overexpressing alongside a RFP-EGFP-LC3 reporter (C57BL/6-Tg(CAG-RFP/EGFP/Map1lc3b)1Hill/J, strain# 027139, The Jackson Laboratory, Bar Harbor, ME USA), as well as their RFP-EGFP-LC3 reporter littermates (8-12 weeks old) were bred and housed in a temperature- and humidity-controlled facility on a 12-h light/dark cycle with free access to food and water. Animals were monitored daily and randomly assigned to experimental groups. Animals that displayed surgical complications or died perioperatively were excluded.

### Murine Model of Myocardial Infarction (MI)

Myocardial ischemia was induced by permanent surgical ligation of the left anterior descending (LAD) coronary artery. Mice were anesthetized with 2% isoflurane delivered in oxygen, intubated with a 20-gauge catheter, and ventilated using a rodent ventilator. A left thoracotomy was performed through the fourth intercostal space to expose the heart. The pericardium was gently opened, and the LAD was visualized running between the pulmonary artery and the left atrium. A 7–0 nylon suture was passed beneath the LAD approximately 1–2 mm from its origin. For permanent MI, the suture was tied securely, resulting in immediate blanching and hypokinesis of the anterior LV wall. Sham-operated mice underwent the same thoracotomy and suture placement without ligation. After closure of the chest wall and extubation, mice were placed on a warming pad until fully recovered.

### RNA Extraction and bulk RNA Sequencing

For transcriptomic profiling, myocardial tissue was harvested from the infarcted left ventricle (LV) of WT and cardiomyocyte-specific FYCO1-transgenic mice 3 days after permanent LAD ligation. Hearts were rapidly excised, and the infarct border zone of the LV was carefully dissected, snap-frozen in liquid nitrogen and stored at −80 °C until further processing. Total RNA was extracted from heart tissue using RNA Extraction Kits (Quick-RNA Miniprep Plus Kit # R1057, ZymoResearch) according to the manufacturer’s instructions. RNA purity and concentration were assessed spectrophotometrically, and RNA integrity was evaluated prior to library preparation. RNA sequencing libraries were generated from high quality total RNA and processed by Novogene (Cambridge, UK) according to the company’s standard workflow. Briefly, mRNA was enriched, fragmented, and reverse transcribed, followed by second strand synthesis, end-pair, adaptor ligation, PCR amplification, and library quality assessment. Sequencing was performed on a Illumina platform to generate paired-end reads. Raw reads underwent quality control, adaptor removal, and filtering of low-quality reads. Clean reads were aligned to mouse reference genome employing a standard splice-aware aligner. Gene expression levels were quantified and normalized, and differential expression analysis between WT and FYCO1-Tg samples was performed using established bioinformatic pipeline. Genes exhibiting adjusted P<0.05 were deemed significantly differentially expressed. Downstream analyses included principal component analyses, heatmap visualization, volcano plot presentation, and gene ontology pathway enrichment.

### Serum cytokine and chemokine profiling

To assess systemic inflammatory responses on protein level following myocardial infarction, blood samples were collected from WT and FYCO1-trangenic mice 3 days post ligation. Whole blood was allowed to clot at room temperature, centrifuged to yield serum, aliquoted, and stored at −80°C until analysis. Circulating levels of cytokine, chemokines and cardiac injury markers were quantified using a multiplex bead-based immunoassay platform (Mouse Cardiovascular Disease Panel 1 7-Plex-Discovery Assay®, Mouse Cardiovascular Disease Panel 2 9-Plex-Discovery Assay®, Mouse Cytokine 44-Plex-Discovery Assay®, Eve Technologies, Calgary, AB, Canada) according to the manufacturer’s instructions. This platform enables simultaneous detection of multiple analytes from small serum volumes using a fluorescent bead-based technology. Assessed analytes included pro-inflammatory cytokines, anti-inflammatory, chemokines, and cardiac Troponin T. Samples were processed in parallel under standardized conditions, with concentrations interpolated from provider-generated standard curves. Data were subsequently used for comparative analysis between WT and FYCO1-Tg groups.

### Western Blotting

Mouse hearts were homogenized in RIPA buffer supplemented with protease and phosphatase inhibitors (complete™, Mini, EDTA-free Protease-Inhibitor-Cocktail # 04693159001, Roche; Phosphatase Inhibitor Cocktail 2, # P5726, Sigma, Phosphatase Inhibitor Cocktail 3, # P0044, Sigma). Protein concentrations were quantified by DC assay. Samples (20–40 μg) were separated by SDS–PAGE, transferred to Nitrocellulose membranes. For quantification of total protein membranes were incubated in Ponceau S (# P7170, Sigma), before being blocked in 5% BSA for 1–2 h. Membranes were incubated overnight at 4°C with antibodies anti-FYCO1 (NovusBiologicals, # 47266), anti-LC3B (CellSignaling, #2775), anti-p62 (CellSignaling, #5114), anti-Atg5 (CellSignaling, #2630), anti-Rab7 (Sigma, # R4779), anti-Bcl-2 (Proteintech, # 68103), anti-Bax (CellSignaling, #14796), Caspase 3 (CellSignaling, #9662), cleaved Caspase 3 (CellSignaling, #9661), Caspase 7 (CellSignaling, #9492), cleaved Caspase 7 (CellSignaling, #9491), Caspase 9 (CellSignaling, #9508), or cleaved Caspase 9 (CellSignaling, #9507). After washing, HRP-conjugated secondary antibodies were applied for 1 h at room temperature, and signal was visualized using enhanced chemiluminescence. Protein targets were normalized against total protein.

### Histology and Immunohistochemistry

Hearts were fixed in 4% paraformaldehyde, dehydrated in a sucrose gradient and embedded in Tissue-Tek O.C.T. (Sakura# 4583). Sections (5 μm) were stained with Masson’s trichrome (Sigma# HT15) according to the manufacturer’s instructions to quantify fibrosis. Immunohistochemistry and immunofluorescence were performed after blocking and incubation with primary antibodies (anti-Mac-2 (Cedarlane, # CL8942AP), anti-CD68 (abcam, ab53444), anti-MCP-1 (abcam, ab308523) or cleaved Caspase 7 (CellSignaling, #9491)). Fluorescent secondary antibodies Alexa Fluor® 647 goat anti-rat (Jackson, # 112-605-167), Alexa Fluor® 546 goat anti-rabbit (Invitrogen, # A-11035) and DAPI were used for visualization by confocal microscopy (Zeiss LSM 880 with Airyscan, Zeiss GmbH, Jena, Germany). Analysis was performed using ImageJ software.

### Assessment of Autophagic Flux (In Vivo Reporter Model)

In vivo, autophagic flux was assessed using RFP-EGFP-LC3 transgenic reporter mice (C57BL/6-Tg(CAG-RFP/EGFP/Map1lc3b)1Hill/J, strain# 027139, The Jackson Laboratory, Bar Harbor, ME USA). Hearts were harvested at designated time points after LAD ligation, fixed in 4% paraformaldehyde, cryoprotected, and sectioned into 5-μm slices. Sections were mounted with antifade medium and imaged by confocal microscopy using identical acquisition settings across groups. Autophagosomes were identified as yellow puncta (EGFP⁺/RFP⁺), whereas autolysosomes were detected as red-only puncta (RFP⁺) due to GFP quenching in the acidic lysosomal environment.

### Infarct Size Measurement (Evans Blue and TTC Staining)

Myocardial area at risk (AAR) and infarct size were assessed 3 days and 30 days post MI using Evans Blue and 2,3,5-triphenyltetrazolium chloride (TTC) staining. Hearts were harvested, and flushed with 1% Evans Blue via retrograde injection through the aorta, while the heart was still beating, to delineate the non-ischemic perfused myocardium, which stained dark blue. Hearts were frozen briefly at −20°C to facilitate slicing.

Transverse sections (1 mm thick) were incubated in 2% TTC at 37°C for 15–20 min. TTC- stained viable myocardium appeared red, whereas infarcted tissue remained pale. The AAR (Evans Blue–negative region), infarct area (TTC-negative region within the AAR), and total left ventricular area were quantified using ImageJ.

### Echocardiography

Cardiac function after myocardial infarction was assessed by transthoracic echocardiography using a high-frequency small-animal imaging system (Vevo 1,100 ultrasound system (VisualSonics, FUJIFILM, Japan)) equipped with a 30–40 MHz transducer. Mice were lightly anesthetized with 1–2% isoflurane to maintain a physiological heart rate, and placed on a temperature-controlled platform. Parasternal long-axis and short-axis views were obtained to visualize the infarcted anterior wall. M- mode recordings at the mid-papillary level were used to measure left ventricular internal diameters at diastole (LVIDd) and systole (LVIDs). Ejection fraction (EF) and fractional shortening (FS) were calculated.

### Statistical Analysis

Data are expressed as mean ± SEM. Comparisons between groups were made by one-way ANOVA followed by Tukey’s multiple-comparison test or by unpaired t-tests where appropriate. Statistical significance was defined as p < 0.05. Analyses were performed using GraphPad Prism (v.11).

## Results

### Cardiomyocyte-specific FYCO1 overexpression limits infarct expansion and preserves contractile function following myocardial infarction

To investigate the potential role of FYCO1 in modulating cardiac injury and remodeling, we subjected cardiomyocyte-specific FYCO1 transgenic (FYCO1-Tg) mice and wild-type (WT) littermates to permanent ligation of the left anterior descending (LAD) coronary artery (Figure 1a). Our previous work demonstrated, that FYCO1 expression was selectively increased in cardiomyocytes of transgenic animals, and baseline cardiac morphology and function were indistinguishable between genotypes [45]. Following permanent coronary artery occlusion, morphology and function were evaluated by echocardiography and histology at 3 and 30 days post ligation. Early injury responses were evaluated 3 days post myocardial infarction (MI). At this time point, FYCO1-Tg mice exhibited significantly reduced levels of circulating Troponin T (Figure 1g), indicating attenuated acute cardiomyocyte injury. Consistent with this observation, histological analysis revealed reduced scar formation (Figure 1f) in transgenic animals compared to WT controls. Despite this reduction in overall tissue injury, reduction in cardiac function was comparable between groups (Figure 1h), as reflected by equivalent ejection fraction (EF) in the early phase post infarction.

**Figure 1:**
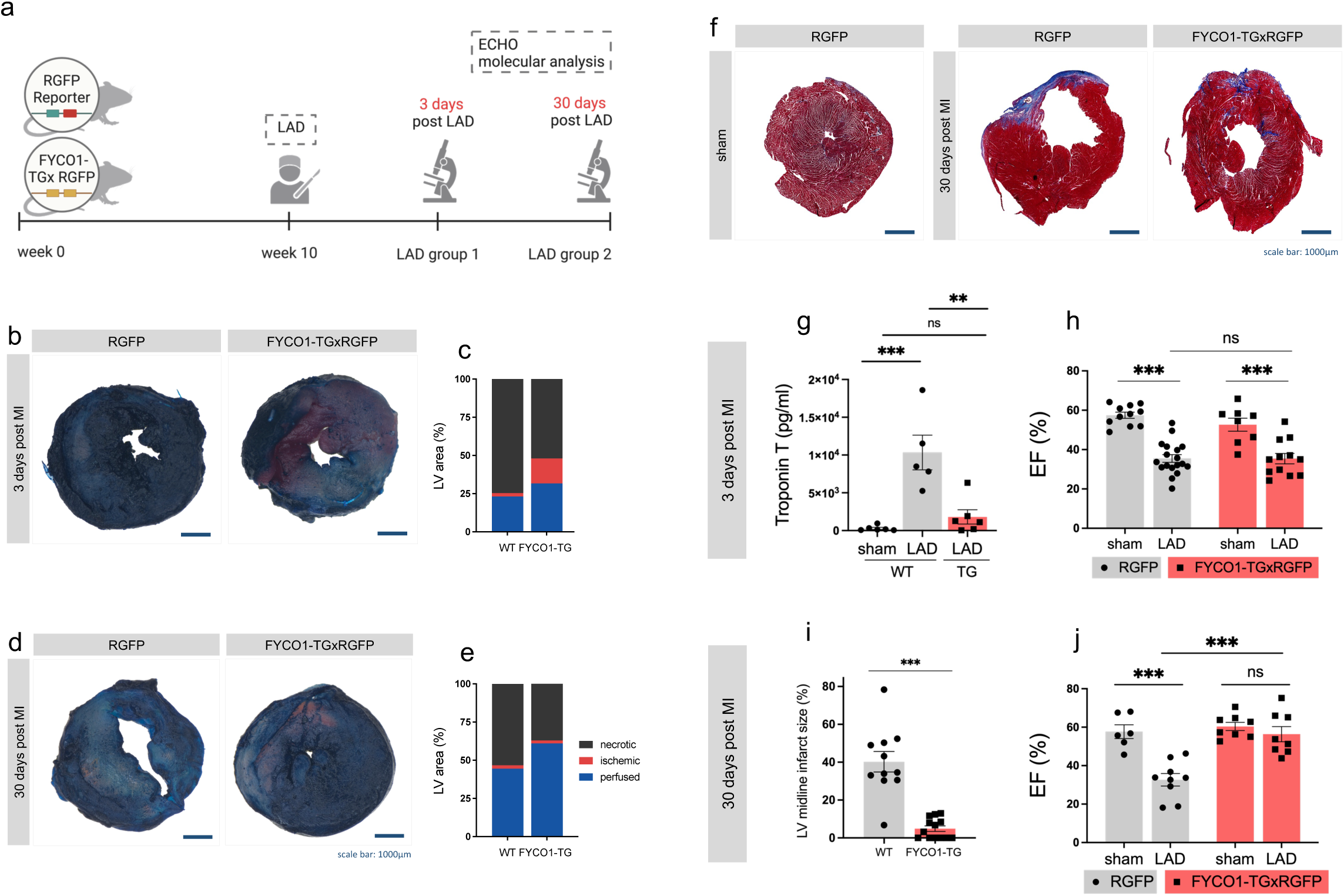
Cardiomyocyte-specific FYCO1 overexpression limits infarct expansion and preserves cardiac function following myocardial infarction. (a) Schematic illustration of the study design of our conducted *in vivo* study. (b,d) Illustrative images of LV sections stained with EvansBlue/TTC-staining 3 days and 30 days post MI. The graphs represent the statistical quantification of perfused, ischemic and necrotic area 3 days (c) and 30 days (e) post MI (n=2-3, 6 sections analyzed per heart sample). (f) Illustrative brightfield images of LV stained with masson trichrome 30 days post MI. (g) Serum Troponin T levels 3 days post MI. Statistical quantification of masson trichrome-stained LV sections for midline infarct size 30 days post MI (i) (n=11- 14, 3 images per specimen). Ejection fraction measurements 3 days (h) and 30 days (j) post MI performed by echocardiography (n=7-18).

We next evaluated the impact of FYCO1 overexpression on long-term post-infarction remodeling. 30 days post MI, when post-infarction remodeling is stable, FYCO1-Tg mice displayed a significantly reduced infarct size (Figure 1i). This structural preservation translated into significant improvement in cardiac function, with transgenic animals displaying near-normal ejection fractions (Figure 1j), while WT mice exhibited persistent contractile impairment.

To further delineate the extent of myocardial injury, we performed Evans Blue/TTC staining to distinguish perfused, ischemic and necrotic myocardial regions.

Interestingly, early after ligation, FYCO1-Tg mice exhibited a larger ischemic but viable myocardial area, suggesting enhanced tissue salvage potential within the area at risk (Figure 1b,c). Consistent with this early-phase observation, analysis after completed remodeling revealed a significantly larger perfused myocardial region in FYCO1-Tg hearts, accompanied by diminished ischemic and necrotic regions compared with WT controls (Figure 1 d,e).

Collectively, these results indicate that cardiomyocyte-specific overexpression mitigates myocardial injury and limits adverse remodeling following MI, ultimately preserving myocardial viability and cardiac function.

### FYCO1 promotes autophagic flux following myocardial infarction

To enable direct in vivo visualization of autophagy dynamics, FYCO1 transgenic mice were crossed with tandem fluorescent RFP-EGFP-LC3 reporter mice [45–47], enabling assessment of autophagosome formation and autolysosomal processing in cardiomyocytes. This reporter expresses autophagosomal marker LC3, fused to two fluorophores with distinct pH sensitivity (Figure 2a). While both green EGFP and red RFP signals are stable in neutral vesicles, EGFP fluorescence is quenched upon lysosomal acidification, allowing for discrimination between autophagosomes and autolysosomes by confocal microscopy. While both fluorophore signals are preserved in autophagosomes, resulting in yellow puncta (RFP+EGFP+; yellow), the acidic environment in autolysosomes quenches EGFP fluorescence, which therefore appear as red puncta (RFP+EGFP-; red). Analysis of confocal microscopy revealed that FYCO1- Tg mice displayed increased numbers of both autophagosomes (RFP+EGFP+; yellow) and autolysosomes (RFP+EGFP-; red) compared to WT mice under basal conditions, indicating enhanced but coordinated and “balanced” aautophagic flux. Specifically, the proportional increase of both vesicles suggest sustained autophagosome formation as well as efficient autolysosomal clearance, indicating an overall augmentation of autophagic flux rather than accumulation of stalled intermediates.

**Figure 2:**
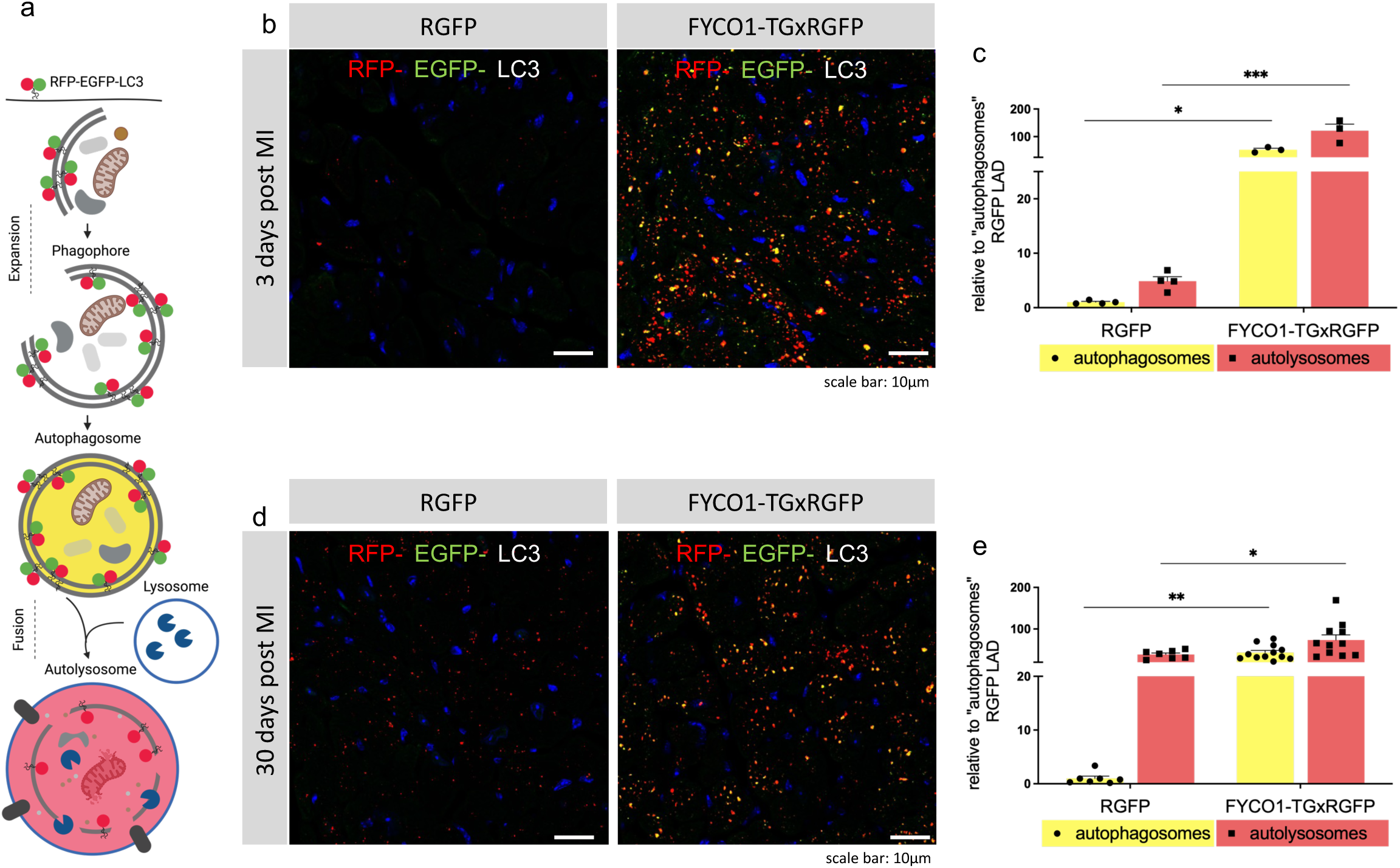
FYCO1 promotes sustained and balanced autophagic flux following myocardial infarction and protects against pathological accumulation of autolysosomes which benefits remodeling. (a) Schematic representation of *in vivo*-RFP-EGFP-LC3-construct. (b,d) Illustrative confocal images showing colocalization of RFP (red) and EGFP (green) of RFP-EGFP-LC3 tandem fluorophore in immunohistochemistry demonstrating autophagosome (RFP+ EGFP+; yellow) and autolysosome (RFP+ EGFP-; red) presence in heart sections of RGFP and FYCO1-TGxRGFP mice. The graphs represent the statistical quantification of colocalization analysis of red and green fluorescence in confocal microscopy images 3 days (c) and 30 days (e) post MI (n=3-9, 6 images analyzed per section).

In WT mice, acute ischemia following ligation induced transient activation of autophagy, consistent with a stress-induced response (Figure 2b,c) [12,14,48]. However, 30 days post MI after completed cardiac remodeling, WT hearts displayed marked accumulation of autolysosomes (Figure 2d,e), suggesting impaired clearance and dysregulated autophagic turnover during cardiac remodeling [49–53]. In contrast, FYCO1-Tg hearts maintained elevated levels of autophagosomes and autolysosomes throughout both the acute and chronic phase after ligation, without evidence of pathological accumulation of either vesicles. Quantitative analysis revealed, that while being overall enhanced – autophagosome formation and autolysosomal clearance remained proportionally coupled, suggesting a sustained autophagic flux. These findings indicate, that FYCO1 enhances the efficiency of the autophagic cascade, promoting continuous cargo processing and preventing maladaptive accumulation of autophagic vesicles observed in WT myocardium (Figure 2d,e).

In WT mice, acute ischemia following ligation induced transient activation of autophagy, consistent with a stress-induced response [12,14,54]. However, 30 days post MI after completed cardiac remodeling, WT hearts displayed marked accumulation of autolysosomes, suggesting impaired clearance and dysregulated autophagic turnover during cardiac remodeling [15,50–53]. In contrast, FYCO1-Tg hearts maintained elevated levels of autophagosomes and autolysosomes throughout both the acute and chronic phase after ligation, without evidence of pathological accumulation of either vesicles. Quantitative analysis revealed, that while being overall enhanced – autophagosome formation and autolysosomal clearance remained proportionally coupled, suggesting a sustained autophagic flux. These findings indicate, that FYCO1 enhances the efficiency of the autophagic cascade, promoting continuous cargo processing and preventing maladaptive accumulation of autophagic vesicles observed in WT myocardium.

### FYCO1 overexpression augments autophagic signaling and autophagosome turnover upon ischemia

To further validate the imaging-based confocal microscopy assessments of increased autophagic flux, we quantified protein expression of key autophagy regulators and markers of autophagosome formation. 3 days post MI, WT mice displayed an anticipated stress-induced activation of autophagy [12,52,55], characterized by increased levels of Atg5, Beclin-1, Rab7, p62, and LC3-II alongside elevated LC3-II/I ratio, consistent with an acute autophagic response to ischemic injury (Figure 3a-l). Conversely, FYCO1-Tg hearts displayed significantly higher levels of these autophagy-related proteins under basal and ischemic conditions, indicating that FYCO1 overexpression not only enhances the last steps of the autophagy cascade but preconditions the entire cardiomyocyte autophagic machinery. Notably, the most prominent differences were observed in LC3-II abundance and the LC3-II/I ratio (Figure 3g,h), canonical indicators of autophagosome formation and turnover. Given that FYCO1 has previously been described in facilitating autophagosomal transport toward lysosomes [45], the substantial increase in LC3-II and LC3-II/I ratio supports our hypothesis that FYCO1 enhances autophagosome formation and efficient flux through the autophagy pathway, while preventing accumulation of vesicles. Consistent with these protein expression data and the assessment of the autophagic flux using the RFP- EGFP-LC3 reporter, we observed a concurrent upregulation of Rab7 (Figure 3l), a key regulator of late endosomal trafficking and autophagosome-lysosome fusion [56,57], again suggesting enhanced FYCO1-dependent progression of autophagosomes toward compartment degradation. Notably, elevated levels of autophagy markers persisted after completion of cardiac remodeling, indicating sustained activation of autophagic machinery in FYCO1-Tg myocardium (Figure 3m-t). In contrast to autophagy dysregulation evident in WT hearts during chronic remodeling, this persistent activation occurred within the context of “balanced” autophagic flux, as supported by results of tandem LC3 reporter confocal analyses (Figure 2d,e). Collectively, these protein expression data support the concept that FYCO1 promotes a sustained and coordinated activation of the autophagy machinery, with efficient autophagosome formation, trafficking and autolysosomal degradation in ischemic myocardium.

**Figure 3:**
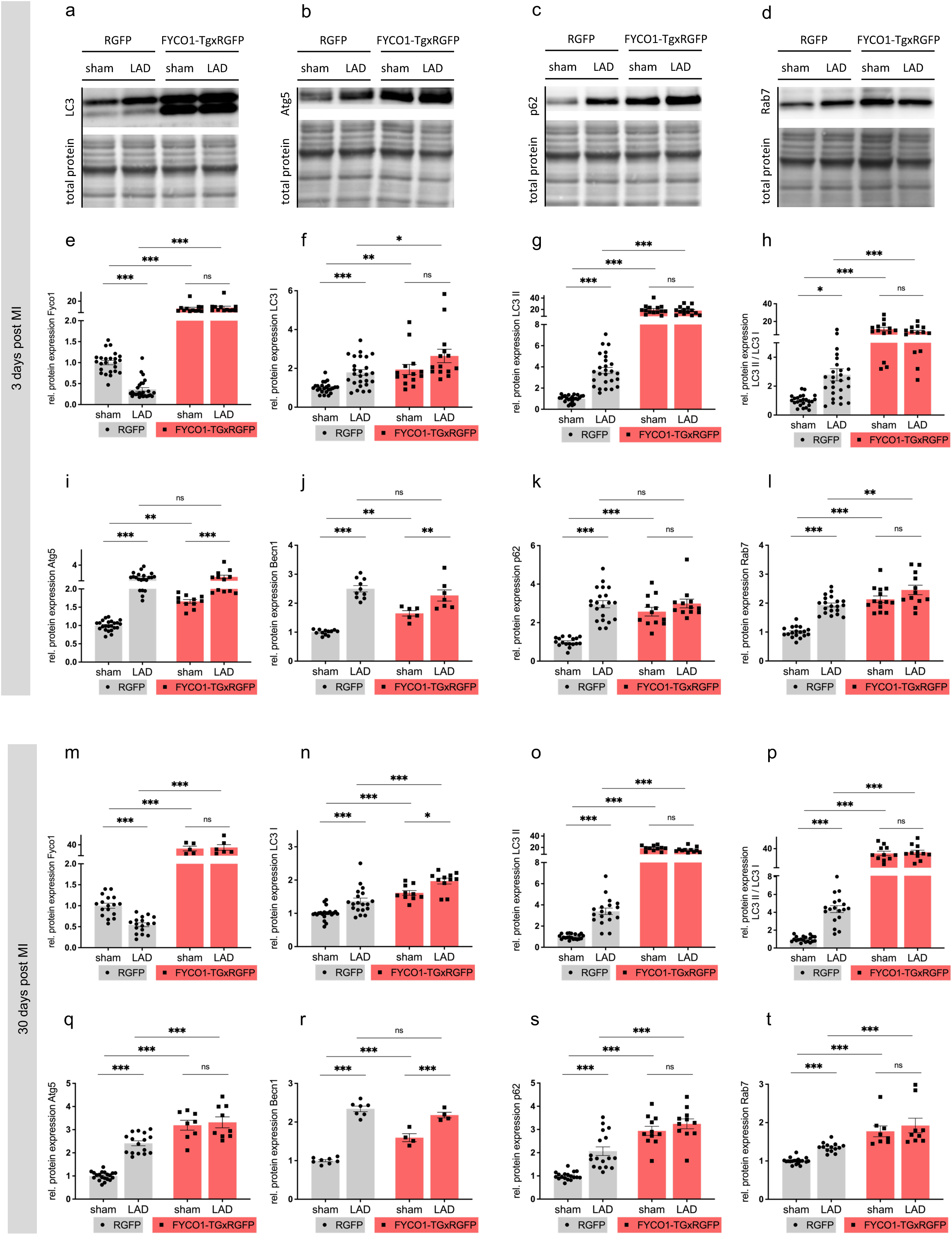
FYCO1 overexpression augments autophagic signaling and autophagosome turnover upon ischemia. (a-d) Westernblot images and quantification of autophagy markers FYCO1 (e, m), LC3 I (f,n), LC3 II (g,o), LC3 II/I (h,p), Atg5 (i,q), Beclin-1 (j,r), p62 (k,s) and Rab7 (l,t) on protein level in FYCO1-TGxRGFP compared to RGFP mice (WT control) 3 days and 30 days post MI. Protein levels determined by densitometry of westernblots, normalized against total protein (n=4-17).

### Transcriptomic profiling reveals a cardioprotective gene program in FYCO1-Tg hearts

To define the molecular programs associated with FYCO1-mediated cardioprotection, we performed bulk RNA sequencing on myocardial tissue from FYCO1-Tg mice and WT controls following ischemic injury. Transcriptomic profiling revealed a distinct gene expression signature in FYCO1-Tg myocardium, characterized by coordinated suppression of inflammatory and pro-apoptotic pathways alongside activation of adaptive tissue remodeling programs. Gene enrichment analysis demonstrated pronounced suppression of pro-inflammatory signaling networks (Figure 4b), including diminished expression of cytokines and chemokines (Figure 4c). In parallel, transcripts associated with apoptotic signaling cascades (Figure 4d) were significantly reduced, indicating blunted cardiomyocyte death programs under FYCO1 overexpression. Notably, several signaling pathways previously implicated in maladaptive cardiac remodeling were also suppressed. In particular, p53 signaling networks, which orchestrate stress responses and cell death following MI, were significantly downregulated in FYCO1-Tg hearts. Transcripts promoting regulated ECM turnover and infarct zone stabilization were upregulated in FYCO1-Tg hearts, while those associated with fibrotic remodeling were attenuated. These transcriptomics shifts align with the observed attenuation of scar size and structural and functional preservation in FYCO1-Tg hearts. Together, these transcriptomic data hints towards a FYCO1-driven shift towards a cardioprotective transcriptional landscape, characterized by inhibition of inflammatory and apoptotic signaling and improved remodeling, potentially supporting functional recovery in response to ischemic injury.

**Figure 4:**
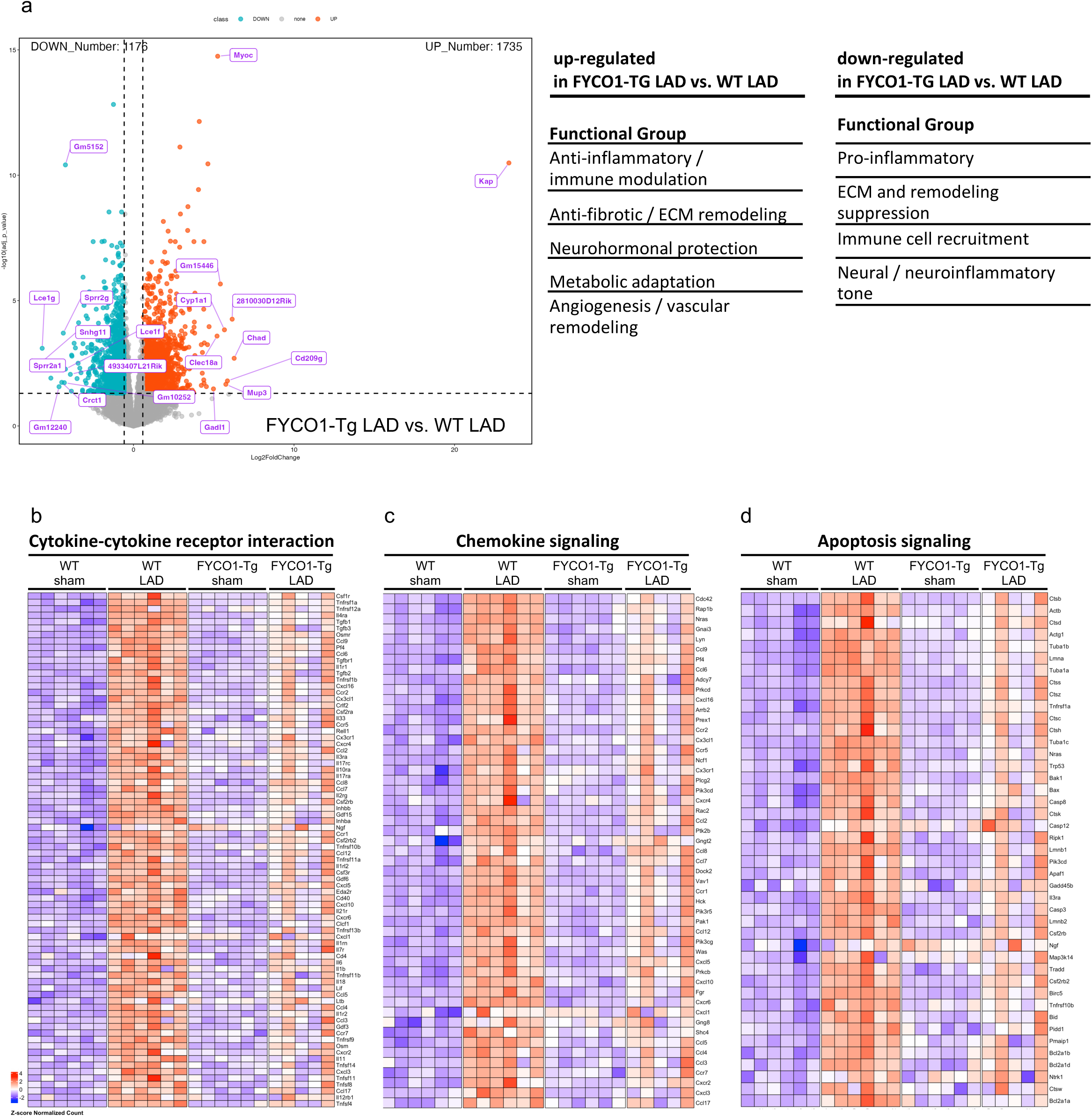
Transcriptomic profiling reveals a cardioprotective gene expression program in FYCO1-Tg hearts. Bulk RNA sequencing was performed on left ventricular tissue obtained from WT and FYCO1-Tg mice 3 days post MI (n=5-6). Differential gene expression analysis revealed a distinct FYCO1-dependent transcriptional signature, characterized by supression of inflammatory and stress-response pathways. (a) Volcano plot showing significantly up- and downregulated genes in FYCO1-Tg LAD versus WT LAD hearts. Heatmaps illustrating differential expression of genes involved in (b) cytokine-cytokine receptor interaction, (c) chemokine signaling and (d) apoptosis signaling.

### Serum cytokine and chemokine screening revealed reduced inflammatory signaling alongside reduced chemoattractant expression upon FYCO1 overexpression

To further delineate the influence of FYCO1 on inflammatory responses following MI, we performed systemic serum cytokine and chemokine profiling in FYCO1-transgenic and WT mice in early phase acute ischemia, using a multiplex bead-based immunoassay platform. Consistent with the cardioprotective transcriptional signatures observed by RNA sequencing, FYCO1-transgenic animals exhibited a significant attenuation of circulating cytokine levels, affecting both pro-inflammatory and anti-inflammatory mediators (Figure 5a). Canonical pro-inflammatory cytokines including IL- 6, IL-1β and INFγ (Figure 5b,c) were significantly diminished in FYCO1-transgenic mice compared to controls. Of note, several cytokines associated with immune cell activation and leukocyte recruitment, such as INFγ and GM-CSF were also found reduced. Interestingly anti-inflammatory cytokines, including IL,10, IL-11 and IL-16 (Figure 5d,e) were also decreased, indicating a global attenuation of systemic inflammation rather than a simple shift toward an anti-inflammatory phenotype. This attenuated anti-inflammatory protein signature translated into an overall reduced chemokine abundance in FYCO1-Tg mice (Figure 5f), particularly for monocyte-attracting chemokines (Figure 5f,g) such as CCL2/MCP-1 and CCL12/MCP-5, indicating reduced recruitment signals for circulating monocytes to injured myocardium. Similarly, several additional chemokines associated with leukocyte migration and immune cell trafficking were decreased, including CCL5/RANTES, CXCL16, CCL3/MIP-1α and CCL21/6Ckine (Figure 5f,h-j), which have been implicated in the recruitment of monocytes, macrophages and T cells during the inflammatory phase following MI [58–60].

**Figure 5:**
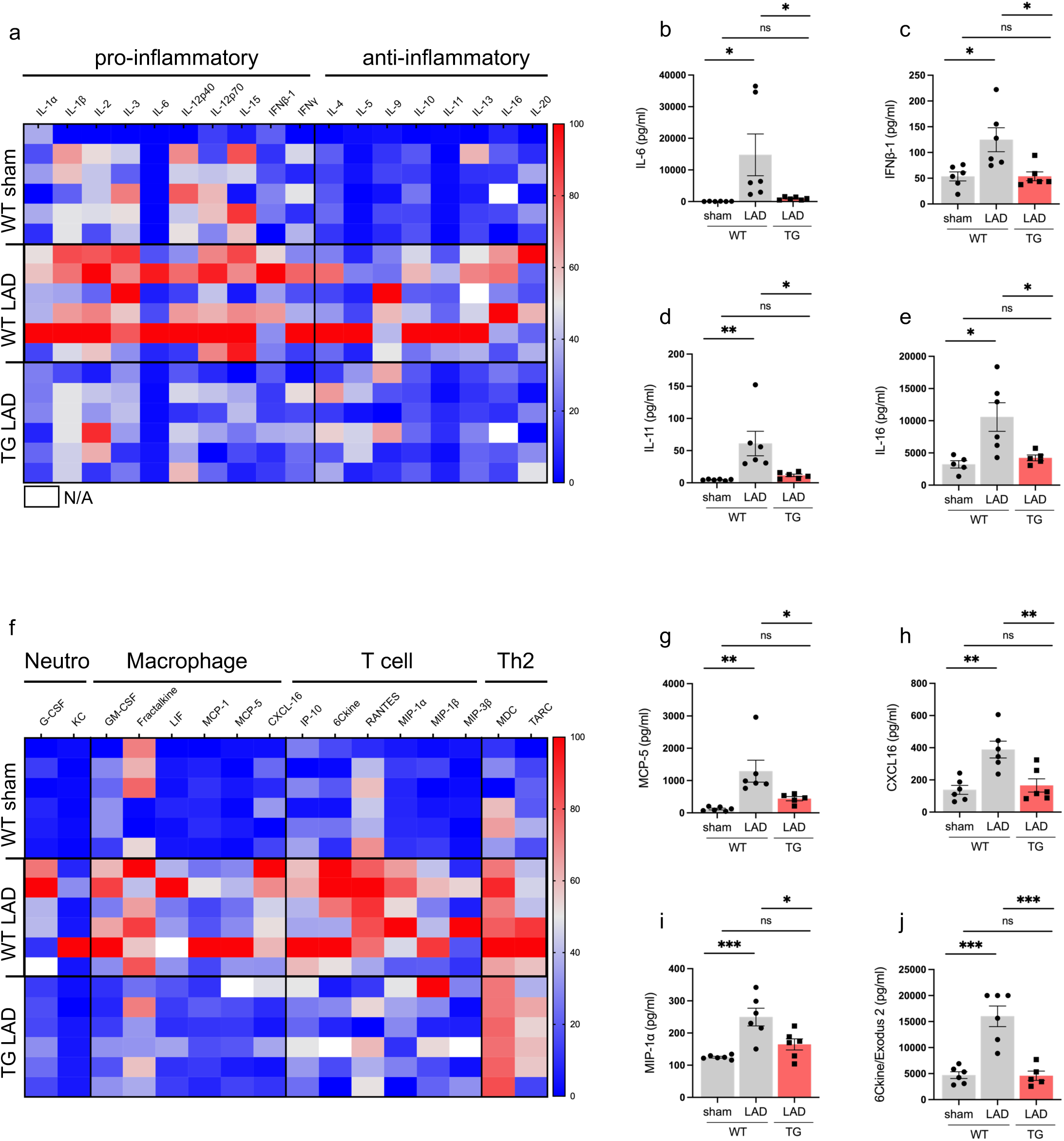
Serum cytokine and chemokine screening revealed reduced inflammatory signaling alongside reduced chemoattractant expression under FYCO1 overexpression. Serum cytokine (a-e) and chemokine (f-j) levels were quantified in WT and FYCO1-Tg mice 3 days following MI (n=6). Quantification of markers was performed using the Mouse Discovery Assay (Eve Technologies, Calgary, AB, Canada), a multiplex bead-based immunoassay platform.

### FYCO1 overexpression limits systemic and local inflammatory signaling and immune cell recruitment into infarct border zone

To determine, whether these systemic signaling patterns translate into altered immune cell recruitment within the myocardium of FYCO1-transgenic mice post MI, we performed immunofluorescence staining of infarct border zone tissue. Macrophage accumulation in the border zone was assessed using the established macrophage markers Mac-2 (Galectin-3) and CD68, in order to identify activated macrophages in infarcted myocardium [58,59]. Confocal imaging revealed a marked reduction of Mac2+ (Figure 6a,b) and CD68+ (Figure 6a,c) macrophages within the infarct border zone of FYCO1-transgenic hearts compared to intense accumulation in WT controls during acute ischemic phase post MI, indicating diminished immune cell recruitment at the site of injury. Consistent with reduced levels of circulating chemokines identified in serum profiling, MCP-1 (CCL2) staining within infarcted myocardium, and known to be localized in injured cardiomyocytes, activated fibroblasts and infiltration immune cells, was profoundly diminished. Monocyte chemoattractant protein MCP-1 is a key chemokine for the recruitment of circulating monocytes during early the inflammatory phase after myocardial infarction [61]. The scarcity of MCP-1 expression in border infarct zone in FYCO1-transgenic myocardium (Figure 6a,d) signifies not only decreased systemic inflammatory signaling but decreased local chemoattractant signaling as well. Quantification of MCP-1 expression in injured myocardium alongside Mac2+ and CD68+ cells across multiple border zone regions confirmed an overall reduction in macrophage recruitment in FYCO1-trangenic hearts. Notably, this reduction of chemoattractant expression and immune cell recruitment occurred despite a comparable initial reduction of ejection fraction, suggesting that FYCO1 actively orchestrates inflammatory signaling following ischemia, and not only being a result of reduced initial myocardial dysfunction.

**Figure 6:**
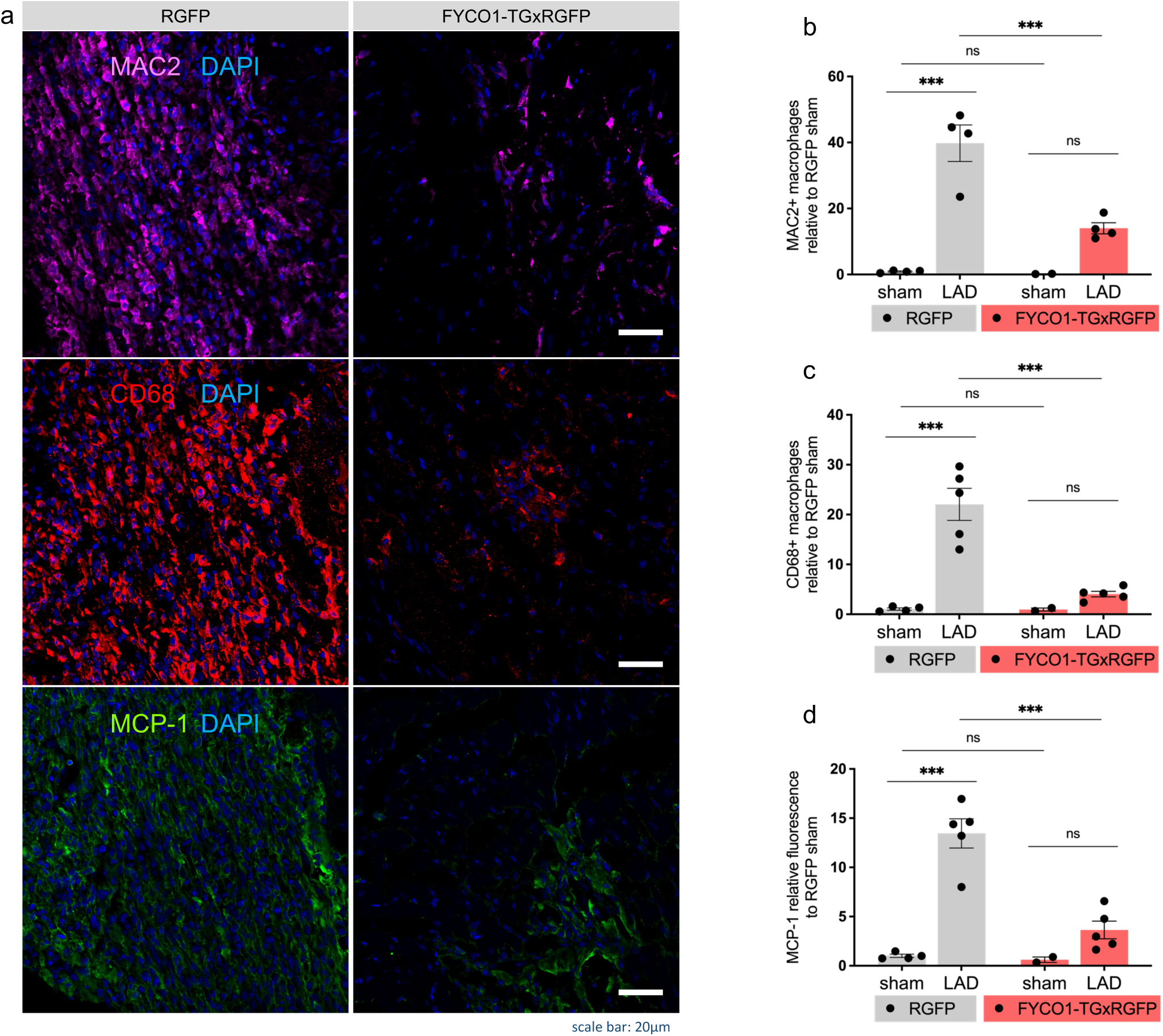
FYCO1 overexpression limits systemic and local inflammatory signaling and immune cell recruitment into infarct border zone. (a) Illustrative confocal images showing macrophage infiltration into the infarct zone 3 days post MI. LV sections are stained with MAC2, CD68 and MCP-1 antibody with DAPI. Graphs represent the statistical quantification of MAC2 (b), CD68 (c) and MCP-1 (d) signal relative to WT control (analyzed with ImageJ; n=2-5, 10 images analyzed per section).

Collectively, these immunofluorescence and serum profiling analyses demonstrate that FYCO1 overexpression supresses both systemic proinflammatory signaling and local chemokine-driven immune cell recruitment within infarct border zone, resulting in a potentially more favourable cardiac microenvironment post injury. This ameliorated inflammatory microenvironment aligns with the observed enhanced myocardial function and improved remodeling (Figure 1) in hearts of FYCO1-transgenic mice.

### FYCO1 overexpression alleviates apoptosis induction in acute ischemia

To further determine how FYCO1-driven autophagy regulation mediates cardioprotection, we next analyzed key regulators and executioners of apoptosis in WT and FYCO1-Tg hearts following injury. Immunoblotting revealed a substantial reduction in the abundance of late executing caspases caspase-3, caspase-7 and caspase-9 in FYCO1-transgenic myocardium compared to WT controls. More importantly, the cleaved (activated) forms of these caspases were markedly decreased, and the ratio of cleaved to full-length caspase proteins was significantly lowered in transgenic hearts. These findings indicate restrained activation of the terminal apoptotic cascade (Figure 7j-o).

**Figure 7:**
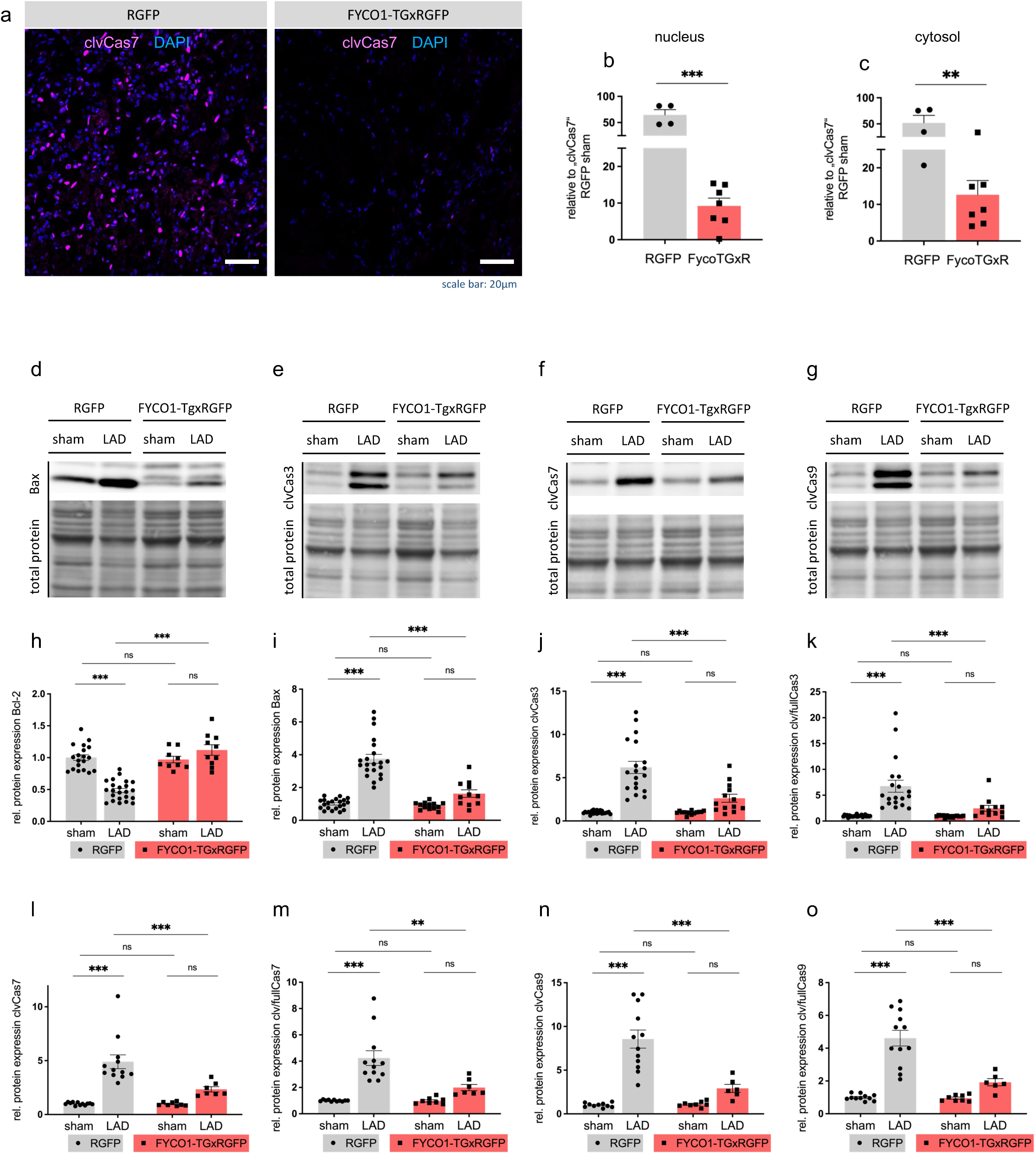
FYCO1 overexpression alleviates apoptosis induction in acute ischemia. (a) Illustrative confocal images showing colocalization of cleaved Caspase7 (pink) and DAPI (blue). Colocalization analysis for the translocalization of cleaved Caspase7 into the nucleus as well as the presence of cleaved Caspase7 in the cytosol. Graphs (b,c) represent the statistical quantification of colocalization analysis of pink and blue fluorescence in confocal microscopy images 3 days days post MI (n=4-7, 12 images analyzed per section). (d-g) Westernblot images and quantification of apoptosis markers Bcl-2 (h), Bax (i), cleaved Caspase 3 (j), cleaved/full Caspase 3 (k), cleaved Caspase 7 (l), cleaved/full Caspase 7 (m), cleaved Caspase 9 (n) and cleaved/full Caspase 9 (o) on protein level in FYCO1-TGxRGFP compared to RGFP mice (WT control) 3 days post MI. Protein levels determined by densitometry of westernblots. (n=6-17).

To spatially delineate apoptotic signaling in the myocardium, we performed confocal immunofluorescence staining for cleaved caspase 7, a pivotal executor of apoptosis during ischemia. In WT hearts, a strong cleaved caspase-7 signal was detected in cardiomyocytes of the infarct border zone. Both cytosolic accumulation and nuclear translocation, hallmarks of active apoptotic cell death execution [62,63], were observed in WT myocardium. In contrast, FYCO1 manifested a scarcity of cleaved caspase-7 positive cells, with significantly diminished signals in both the cytosol and the nucleus, signifying alleviated activation and propagation of apoptosis signaling in cardiomyocytes of these mice.

Consistent with suppression of the late steps in the apoptotic cascade, analysis of upstream mitochondrial-induced apoptosis regulators further supported an anti-apoptotic and cardioprotective microenvironment in FYCO1-Tg myocardium. The pro-apoptotic Bcl-2 family member Bax (Figure 7i) was significantly downregulated, while the the anti-apoptotic protein Bcl-2 (Figure 7h) was upregulated in transgenic hearts. This shift in the Bax/Bcl-2 balance towards a survival-promoting state suggests a FYCO1-dependent stabilization of mitochondrial integrity, which reduces upstream activation of intrinsic apoptotic pathways as well as augmentation of proinflammatory signaling.

Collectively, these findings demonstrate that FYCO1 overexpression suppresses activation of both upstream mitochondrial-derived regulators and downstream execution components of the apoptotic machinery, resulting in reduced apoptotic cell death following ischemia. This attenuation of apoptotic signaling, orchestrated by FYCO1-driven autophagy activation, likely contributes to the improved myocardial integrity and functional preservation observed in FYCO1-Tg mice following myocardial infarction.

## Discussion

Here, we demonstrate that overexpression of FYCO1 robustly enhances autophagic flux in the ischemic myocardium, leading to attenuation of cardiac injury, inflammatory burden, and post-infarct remodeling. Moreover, we provide mechanistic evidence that restoring autophagic flux—rather than simply stimulating autophagosome formation— is a key determinant of cardiomyocyte survival after acute myocardial infarction. As impaired autophagosome–lysosome fusion and defective degradation have been shown to be central contributors to ischemia–reperfusion injury [12,15,50,56,64], these data position FYCO1 as a potentially targetable node in myocardial protection signaling.

We show that enhancing autophagic flux, rather than merely increasing autophagosome number, is beneficial for protecting the ischemic myocardium. While ischemia is known to transiently induce autophagy as a stress response, inefficient autophagic processing and impaired lysosomal clearance frequently occur during remodeling, resulting in accumulation of dysfunctional proteins and organelles that exacerbate adverse remodeling [15,65]. Several clinical and translational studies have described increased LC3-II and Beclin-1 levels during MI or in failing human hearts [12,19,55,66], but these changes often coexist with p62 accumulation [15,19,67,68], suggesting impaired clearance rather than efficient turnover [14,64]. This distinction is critical: Accumulated autophagosomes without lysosomal degradation or accumulation of autolysosomes may worsen proteotoxic stress, fuel mitochondrial dysfunction, and promote apoptosis [12,53,54,57,64,69]. In contrast, utilizing a RFP-EGFP-LC3 in vivo reporter [45–47], our results show that FYCO1 overexpression enhances autophagic flux, likely by facilitating autophagosomal transport alongside promoting lysosomal degradation, thereby maintaining efficient cargo degradation throughout both acute and chronic phases following myocardial infarction. Specifically, we observed enhanced levels of both autophagosomes and autolysosomes and the absence of accumulation of either vesicles, consistent with a robust and balanced increase in autophagic flux.

This enhancement of autophagic flux in FYCO1-overexpressing hearts was accompanied by increased levels of several other key autophagy markers—including Atg5, p62, Beclin-1 and Rab7. This sustained increased autophagic flux appears critical for preserving cellular homeostasis within the ischemic myocardium, promoting enhanced clearance of damaged components and preventing maladaptive remodeling. This integrates well with prior studies suggesting that lysosomal dysfunction represents a rate-limiting step in autophagy during ischemia–reperfusion injury [12,50,56], as observed in lysosomal storage diseases where defective autophagosome–lysosome fusion leads to autophagosome accumulation and cardiomyopathy [12,51–53,57,69–74], and that interventions targeting this bottleneck—such as spermidine treatment to enhance lysosomal function and flux—confer significant cardioprotection [75,76].

Transcriptomic and serum cytokine and chemokine profiling analyses indicate that FYCO1-mediated autophagy profoundly reshapes the inflammatory landscape within the injured myocardium. Myocardial infarction triggers a tightly coordinated inflammatory response, wherein damaged cardiomyocytes release danger-associated molecular patterns that provoke cytokine and chemokine secretion and drive rapid recruitment of circulating immune cells into the infarct border zone [4,31,33,34]. This early inflammatory phase is dominated by proinflammatory monocytes and macrophages, which collectively participate in clearance of necrotic debris thereby modulating myocardial remodeling [4,33,60,77,78]. However, excessive inflammation early post-MI contributes to cardiomyocyte death, extracellular matrix degradation, and adverse remodeling [9,11,31,41]. While a certain amount of inflammation is required for debris clearance and wound healing, persistent or excessive activation of innate immunity exacerbates myocardial damage [4,58,74,79,80].

Autophagy plays a central role in controlling innate immune activation by degrading damaged mitochondria, aggregated proteins, and intracellular DAMPs before they can trigger downstream inflammatory signaling [12,28,81]. Previous studies have demonstrated that impaired autophagic flux amplifies NLRP3 activation and promotes cytokine secretion, exacerbating post-ischemic tissue injury [64,82]. By restoring lysosomal degradation and reducing substrate overload, these upstream triggers are attenuated, thereby curtailing inflammasome-driven cytokine production. In this context, an interesting and unexpected finding is that FYCO1 overexpression attenuates pro-inflammatory signaling and immune cell infiltration into the infarct border zone.

Our data show significant reductions in pro-inflammatory cytokines, and other pro-inflammatory mediators as well as circulating chemokines such as CCL2/MCP-1, CCL12/MCP-5, CCL5/RANTES, CXCL16, CCL3/MIP-1α and CCL21/6Ckine in FYCO1- overexpressing mice, accompanied by markedly diminished recruitment of neutrophils, monocytes, and macrophages to the infarct border zone [4,11]. Reduced levels of circulating and tissue chemokines—including CCL2/MCP-1, a key regulator of monocyte mobilization from the bone marrow and splenic reservoirs—may underlie the decreased presence of inflammatory macrophages in the border zone [83,84]. Previous studies have shown that CCL2/MCP-1 signaling is a major determinant of monocyte accumulation and macrophage-driven matrix degradation after MI [61,84]. Inhibition of this axis reduces infarct expansion and supports adaptive healing. Consistently, immunofluorescence analysis of FYCO1-Tg myocardium revealed lower MCP-1 expression in cardiac tissue, as well as a decrease in total infiltrating macrophages and a shift away from pro-inflammatory macrophage phenotypes. Moreover, our data indicate not only a reduction in total macrophages but also a shift in macrophage polarization, with fewer pro-inflammatory macrophages present in FYCO1- overexpressing hearts. Excessive M1 macrophage activity has been associated with matrix degradation, heightened oxidative stress, and impaired angiogenesis during post-MI remodeling [58,84]. In contrast, a balanced or M2-skewing macrophage response supports debris clearance, tissue repair, and scar maturation [29,59,60,77,84–86]. Together, these findings indicate that FYCO1 alters the early post-MI inflammatory milieu, creating a myocardial environment more conducive to cardiomyocyte survival and functional recovery. We hypothesize that the anti-inflammatory actions of FYCO1 are likely rooted in its ability to maintain intact and appropriately regulated autophagic flux, thereby promoting removal of defective organelles or other cellular debris such as defective proteins.

Our data further show that enhanced autophagic flux attenuates apoptosis induction in the infarct border zone. Specifically, in FYCO1-overexpressing mice, we observed a significant reduction of protein levels of both key executioner and initiator components of the apoptotic cascade, including Bax, cleaved caspase-3, caspase-7, and caspase-9.

At the same time, expression of the anti-apoptotic protein Bcl-2 was preserved. Likewise, immunofluorescence staining revealed a pronounced decrease in nuclear cleaved caspase-7 signal, indicating that not only activation but also nuclear translocation of executioner caspase-7 is limited in FYCO1-overexpressing hearts. Collectively, these findings identify FYCO1 as a potent modulator of the balance between survival and death signaling in cardiomyocytes exposed to ischemic stress. These observations are in line with the well-established crosstalk between autophagy and apoptosis, particularly via the Beclin-1–Bcl-2 interaction and shared stress pathways such as JNK and p38 MAPK [87–92]. When autophagic flux is impaired, damaged mitochondria accumulate and become a major source of cytochrome c release, subsequent caspase-9 activation, and downstream caspase-3/7–mediated execution [12,64]. Several studies in myocardial ischemia–reperfusion models have shown that blocking autophagy or disrupting mitophagy exacerbates ROS production and apoptotic cell death, whereas interventions that restore autophagic flux blunt caspase activation and reduce infarct size [12,64,93–95]. In this context, our finding that FYCO1 enhances autophagic clearance and simultaneously reduces caspase-3, caspase-7, and caspase-9 activation supports a model in which efficient removal of dysfunctional organelles is a key mechanism underlying its anti-apoptotic effects.

In parallel, improved protein quality control via autophagy-mediated homeostasis preserves ATP production under ischemic conditions and thus confers protection against apoptosis as well as a shift of cell death toward necrosis and necroptosis [12,50,75,64,70,96]. Importantly, apoptotic loss of cardiomyocytes is a major determinant of infarct expansion, wall thinning, and adverse ventricular remodeling in both experimental models and patients [11,97–99]. Thus, enhanced autophagic flux via FYCO1 may protect cardiomyocytes on multiple levels: it removes damaged organelles before they can initiate death signaling, prevents an excessive inflammatory response, and maintains a more favourable energetic state [12,27,52,64,73,100–104]. Prior work has shown that even modest reductions in cardiomyocyte apoptosis after MI translate into meaningful improvements in left ventricular function and long-term outcomes [97–99]. In our study, the combination of diminished nuclear cleaved caspase-7 and lower levels of caspase-3/7/9 is accompanied by a higher ejection fraction, smaller infarct size, and less fibrosis in FYCO1-overexpressing mice. These associations suggest that the anti-apoptotic actions of FYCO1 contribute to an improved structural and functional phenotype post MI.

FYCO1 overexpression preserved left ventricular ejection fraction, reduced infarct size, and attenuated adverse ventricular dilation. These improvements in cardiac function likely stem from the combined effects described above: reduced apoptosis, diminished inflammatory injury, and enhanced clearance of damaged organelles. Structurally, Masson’s trichrome staining revealed decreased fibrosis and improved scar organization in FYCO1–overexpressing mice. Excessive fibrosis is a hallmark of maladaptive remodeling that stiffens the ventricle and impairs systolic and diastolic function [4,5,11]. By preserving viable myocardium and reducing inflammatory and apoptotic signaling, FYCO1 mitigates the stimuli that drive collagen deposition. These findings align with prior translational evidence demonstrating that interventions improving autophagic flux—whether through metabolic stimuli [12,55,70], genetic modulation [12,14,45,57,98], or pharmacologic agents [12,24,25,49,76,105,106]—may favourably influence post-MI remodeling [15,56,107–109].

### Translational Perspectives

Despite accumulating mechanistic evidence supporting a protective role of autophagy in MI, no clinically approved therapies exist that specifically target autophagic flux [20,73,110]. Most candidate autophagy-enhancing interventions in humans exert broad, pleiotropic metabolic effects and do not directly modulate autophagosome– lysosome fusion or lysosomal clearance [73,108,111,112]. Early-phase trials of trehalose [48,107,113], spermidine [76,106,114–117], and KH-1 [118] demonstrate feasibility but remain largely exploratory and have not demonstrated targeted flux correction. Our findings suggest that FYCO1, which improves autophagic flux in a targeted manner and yields substantial cardioprotection in vivo, may represent another target potentially of interest for future translational studies. Additional work could aim to identify upstream activators of FYCO1, develop small molecules that stabilize or enhance its activity, or use peptide mimetics that preserve its functional domains.

### Limitations and Future Directions

Although our study provides mechanistic and functional evidence for FYCO1-mediated cardioprotection, several limitations deserve to be mentioned. First, the use of overexpression models may not reflect the physiological regulation of FYCO1; dose– response studies and loss-of-function experiments will thus be necessary to define the boundaries of its protective effects. Second, as autophagic flux influences many cellular processes, some observed benefits—such as reduced inflammation—may arise from secondary or downstream effects. Parsing direct versus indirect contributions of FYCO1 will require more targeted pathway analysis and the analysis of communication pathways between cell types, alongside its manipulation. Finally, translation to human disease requires a deeper understanding of FYCO1 expression patterns in human myocardium, particularly in patient cohorts exhibiting impaired myocardial autophagic flux. Integration with clinical biomarker data could help to identify patients who might benefit from potential flux-targeting therapies.

### Conclusions and outlook

Collectively, our findings suggest that FYCO1, via enhancing autophagic flux, improves cardiomyocyte homeostasis upon ischemia and modulates the post-MI microenvironment from one dominated by inflammatory injury and cell death towards cell survival, repair, and improved remodeling. This combined mechanism— suppression of inflammatory signaling and apoptotic cell death via improved autophagic flux and likely indirect reduction of immune recruitment through decreased cardiomyocyte death and governed mitochondria quality control—may provide a coherent biological explanation for the robust cardioprotective effects observed. Our results align with emerging literature demonstrating that therapeutic strategies enhancing autophagic flux, can modulate the inflammatory milieu and improve myocardial healing. Our study extends this concept by identifying FYCO1 as a specific regulator that integrates autophagy, inflammation, apoptotic cell death and cell survival towards a cardioprotective response.

## Acknowledgements

F.S. conceptualized the study, designed and performed most of the experiments, analyzed data and wrote the first draft of the manuscript. S.S.H., A.R., T.B., and N.S. performed experiments. C.K., A.M.G and A.Y.R. analyzed data. O.J.M., A.B. and J.H. supervised experiments and critically revised the manuscript. N.F. initiated, designed and conceptualized the study, supervised experiments and prepared the final draft of the manuscript. All authors read and approved the manuscript.

## Sources of Funding

This study was supported by SFB1550-project ID464424253: Collaborative Research Center 1550 (CRC1550) “Molecular Circuits of Heart Disease” projects B04 (to A.B.), B01 (to J.H.), and B03 (to N.F.) by the Deutsche Forschungsgemeinschaft.

## Disclosures

None.

## Nonstandard Abbreviations and Aconyms

6Ckine: C-C motif chemokine ligand 21
AAR: Area at risk
AMI: Acute myocardial infarction
AMPK: AMP-activated protein kinase
ATG5: Autophagy related protein 5
Bax: Bcl-2-associated X protein
Bcl-2: B-cell lymphoma 2
BSA: Bovine serum albumin
CCL2: C-C motif chemokine ligand 2
CD68: Cluster of Differentiation 68
CXCL16: C-X-C motif chemokine ligand 16
DAMPS: Danger-associated molecular patterns
DC: Detergent compatible
EF: Ejection fraction
e.g.: exempli gratia
EGFP: Enhanced Green Fluorescent Protein
FS: Fractional shortening
FYCO1: FYVE and coiled-coil domain autophagy adaptor 1
GLP-1: Glucagon-like Peptide-1
GM-CSF: Granulocyte-Macrophage Colony-Stimulating Factor
HRP: Horseradish peroxidase
INF: Interferon
IL: Interleukin
JNK: c-Jun N-terminal kinase
KH-1: poly(C)-binding protein
LAD: left anterior descending artery
LC3: Microtubule-associated protein 1 light chain 3
LV: Left ventricle
LVIDd: left ventricular internal diameters at diastole
LVIDs: left ventricular internal diameters at systole
Mac-2: Galectin-3
MCP-1: Monocyte chemoattractant protein-1
MI: Myocardial infarction
MIP1α: Macrophage Inflammatory Protein-1 alpha
mRNA: Messenger Ribonucleic acid
NLRP3: NLR family pyrin domain containing 3
P38 MAPK: p38 mitogen-activated protein kinase
p53: Tumor protein p53
P62/SQSTM1: Sequestosome-1
PCR: Polymerase Chain Reaction
Rab7: RAB7, member RAS oncogene family
RANTES: Regulated on Activation, Normal T cell Expressed and Secreted
RFP: Red Fluorescent Protein
RIPA: Radio-Immunoprecipitation Assay
RNA: Ribonucleic acid
SDS-PAGE: Sodium Dodecyl Sulfate-Polyacrylamide Gel Electrophoresis
SEM: Standard Error of the Mean
Tg: transgenic
TTC: 2,3,5-triphenyltetrazolium chloride
WT: Wild-type

## Notes

### Competing Interest Statement

The authors have declared no competing interest.

### Summary of Updates

I just needed to change the middle name of one Co-Author. Nothing else was changed.

